# Characteristics of DNA methylation and gene expression in regulatory features on the Infinium 450k Beadchip

**DOI:** 10.1101/032862

**Authors:** David Martino, Richard Saffery

**Affiliations:** Murdoch Childrens Research Institute, Melbourne, 3052, Australia; University of Melbourne, Melbourne, 3052, Australia

## Abstract

Understanding the relationship between variations in DNA methylation and gene expression has been challenging. Evidence suggests the function of DNA methylation may vary with genomic context, and few consistent rules linking methylation to expression have been noted. For array-based studies, the content of current DNA methylation array platforms provide broad coverage of the genome but target only a fraction of the potentially methylated CG dinucleotides. A better understanding of the interplay between DNA methylation and gene expression is beneficial for users of these platforms, and may aid with candidate prioritization in epigenome-wide association studies (EWAS). To address this we examined the relationship between DNA methylation levels and gene expression in primary T-lymphocytes at discreet genomic regions around the transcriptional unit (Promoters, gene body, untranslated regions) and at CpG island-associated regions (islands, shores and shelves), stratifying by high and low expressed genes. As anticipated we found evidence that DNA methylation at CpG sites near promoter regions are tightly correlated with gene expression in both the stably expressed and developmentally regulated genes, however this is dependent on CpG density. DNA methylation within the gene body was not consistently associated with changes in gene expression. CpG islands and island shores exhibited strong correlations with gene expression, but this was not true for island shelves. We found these relationships were generally preserved at both dynamic and steady state genes, with some notable exceptions. In combination these insights may be useful for prioritising candidates identified in epigenome-wide association studies for subsequent functional studies.

## Introduction

DNA methylation was initially described as an epigenetic gene silencing mark several decades ago,^1^ however since the advent of genome-scale technologies this traditional view has now been expanded. The emerging picture suggests the function of DNA methylation may vary depending upon the context, with evidence to suggest the position of DNA methylation relative to the transcriptional unit may affect gene activity in differing ways.^2^ DNA methylation at the transcriptional start site (TSS) inhibits the binding of DNA polymerase or transcription factors, and hence gene transcription^3^.^4^ The role of DNA methylation within the gene body is poorly understood,^5^ with evidence for both positive and negative correlations with transcription.^6^ These relationships also often vary depending upon underlying CpG density (CpG islands) in a tissue-specific, and developmental manner.

An increasing number of epigenome-wide association studies (EWAS) have been reported in recent years spanning cancer research, epigenetic epidemiology and other complex diseases. The major platform utilized in these studies is the Infinium HumanMethylation450 bead array that measures DNA methylation at over 480,000 genomic CpG sites, and has now been superseded by the Infinium MethylationEpic bead array, which covers over 850 000 methylation sites per sample. These platforms are favoured for large cohort-based studies due to their cost-effectiveness and high sample throughput. A typical association study might yield many potential candidate loci associated with disease or exposure, yet there is little agreement over how to prioritise these for replication and downstream functional studies. In the setting of epigenetic epidemiology, researchers are often limited to using heterogeneous tissues such as blood in cohort-based studies where effect sizes are notoriously small (within the order of 1-2 % differential methylation) and of unknown biological significance. Whilst we are cognizant that the role of DNA methylation may extend beyond regulating gene transcription, researchers are often interested in the likely consequences for gene expression and often direct examination of genome-wide expression is required to assess functional relevance. The Infinium methyaltion BeadChips contain features that are annotated at various regions around gene sequences including promoters, untranslated regions and gene bodies. Here we directly examined the relationship between CpG methylation and gene expression at with a view to understanding which of these annotated regions are more likely to be clearly associated with the regulation of gene expression. We focused on these sites as potentially relevant for changes in gene expression, whilst intragenic regions and enhancer elements are less well defined and more likely to mediate tissue specific effects. We used matched DNA methylation and gene expression measures from publicly available data generated by us previously.^7^ Briefly, DNA and mRNA were co-purified from CD4+ T-cells collected at birth and 12-months from the same volunteers. DNA methylation measures were derived from the Infinium HumanMethylation450 (HM450k) bead array, and gene expression was measured on the Affymetrix Human Gene 1.0ST arrays. Our observations are not intended to be an exhaustive study of the relationship between methylation and expression, instead providing a broad-based view of this relationship to enable the prioritization of loci generated in EWAS analyses for users of these platforms.

## Results

To evaluate the relationship between methylation and gene expression we divided the distribution of gene expression measurements across all samples into quartiles, with the upper and lower quartiles used to define genes with on average low versus high expression respectively. We then compared the kernel density distributions (methylation distributions) of all CpG localised to highly expressed genes, versus CpG localized to genes of low expression according using genomic coordinates provided by the IlluminaHumanMethylation450k.db package. The density distributions were weighted by the number of observations in each class of high and low expression so as to be directly comparable. As shown in figure 1, CpG localised to highly expressed genes have a substantially larger unmethylated area of the beta distribution, where methylation levels range between 0-0.2 (0-20 % methylated), whilst CpG localized to genes of low expression exhibit methylation values that range between 0.8-1.0 (80-100 % methylated) (Figure 1). This anti-correlated relationship is consistent with a generalized classical gene silencing effect overall (i.e. across all genomic regions) but there are several CpG site that do not fit this relationship, being highly methylated yet mapping to highly expressed genes, and vice versa. To explore this further we repeated this analysis stratified by genomic region.

**Figure 1.**
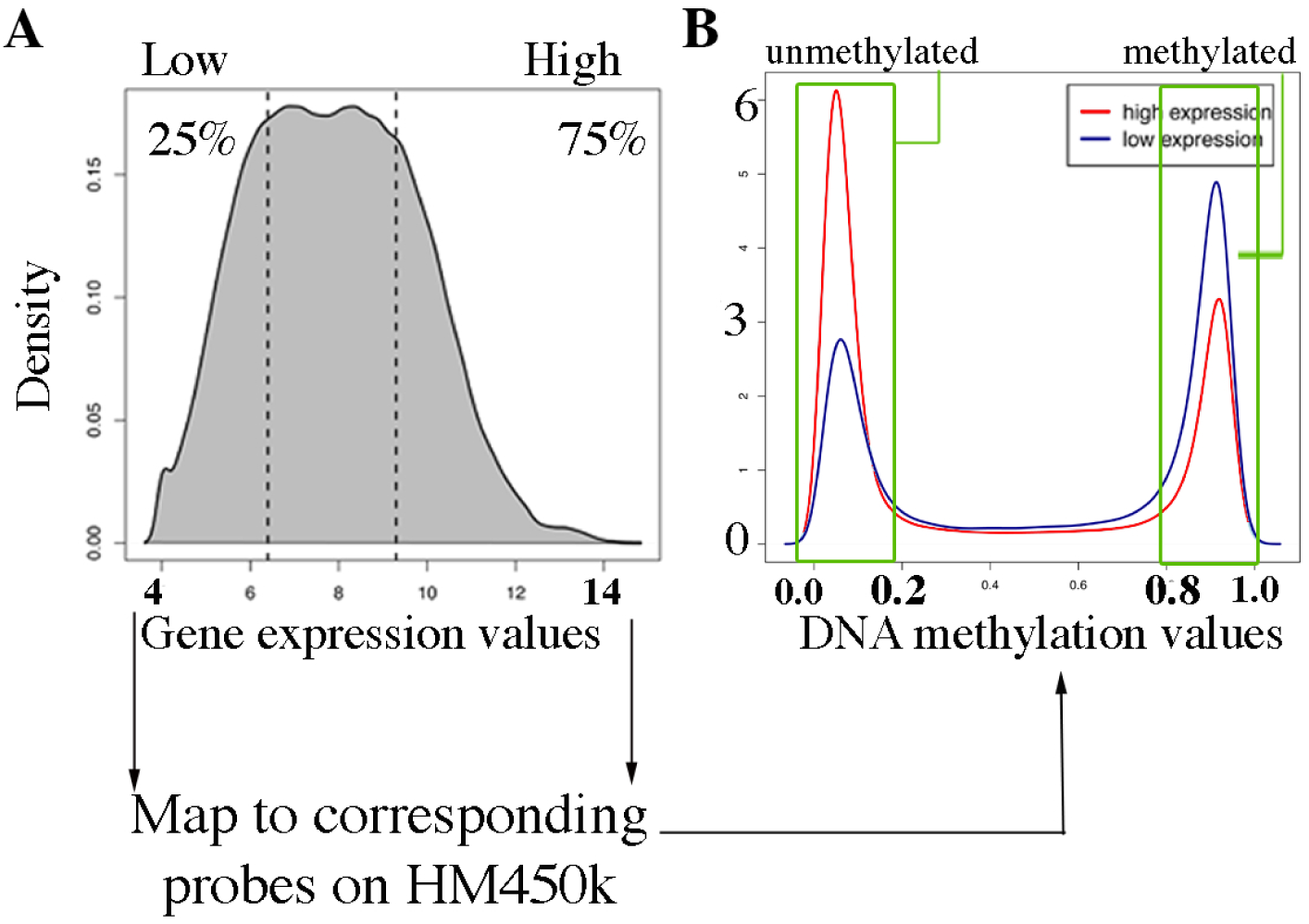
Summary of the methodology. (A) Genes with high versus low expression were defined according to the 25 percent and 75 percent quartiles of the distribution of gene expression measurements. (B) These genes were separately mapped to corresponding probes on the HM450k array by genomic coordinates and the density distribution of DNA methylation measurements associated with these genes was compared. Green boxes indicate the area under the curve for the highly methylated and lowly methylated CpG sites corresponding to these genes. Data shown here represents all genomic sub-regions. The difference between high and low expressed genes is visible as a difference in the peak height ratio of methylated to unmethylated CpG sites.

Next we examined these relationships more specifically at genomic regions around the transcription unit. We noted a range of differences in the relationship between DNA methylation and expression for probes localised across the transcriptional start site (TSS). At close proximity to the promoter (within 200bp of TSS) and within the 1st exon, the relationship between DNA methylation and gene expression was generally inversely correlated as anticipated, consistent with a role for gene silencing in these regions (TSS200 and 1st exon, figure 2A boxes in red). At more distal 5’ UTR regions or 1500bp upstream of the promoter this relationship is less clear (TSS1500, 5’UTR). Interestingly regardless of gene expression level, CpG sites in the gene body or at the 3’UTR are generally highly methylated (figure 2A, boxes in green). At high-density CpG promoter regions (those that overlap CpG islands), CpG sites are highly likely to be hypomethylated, and the level of de-methylation may discriminate gene expression level, with the exception of CpG sites within the 3’UTR (figure 2A boxes in purple). The distribution of DNA methylation sites for high and low expressed genes found at low CpG density 3’UTR regions were not significantly different by the Wilcoxon test, suggesting a lack of direct relationship between methylation status and gene expression in this region. These realtionships held true for both birth and 12-month samples (data not shown).

**Figure 2.**
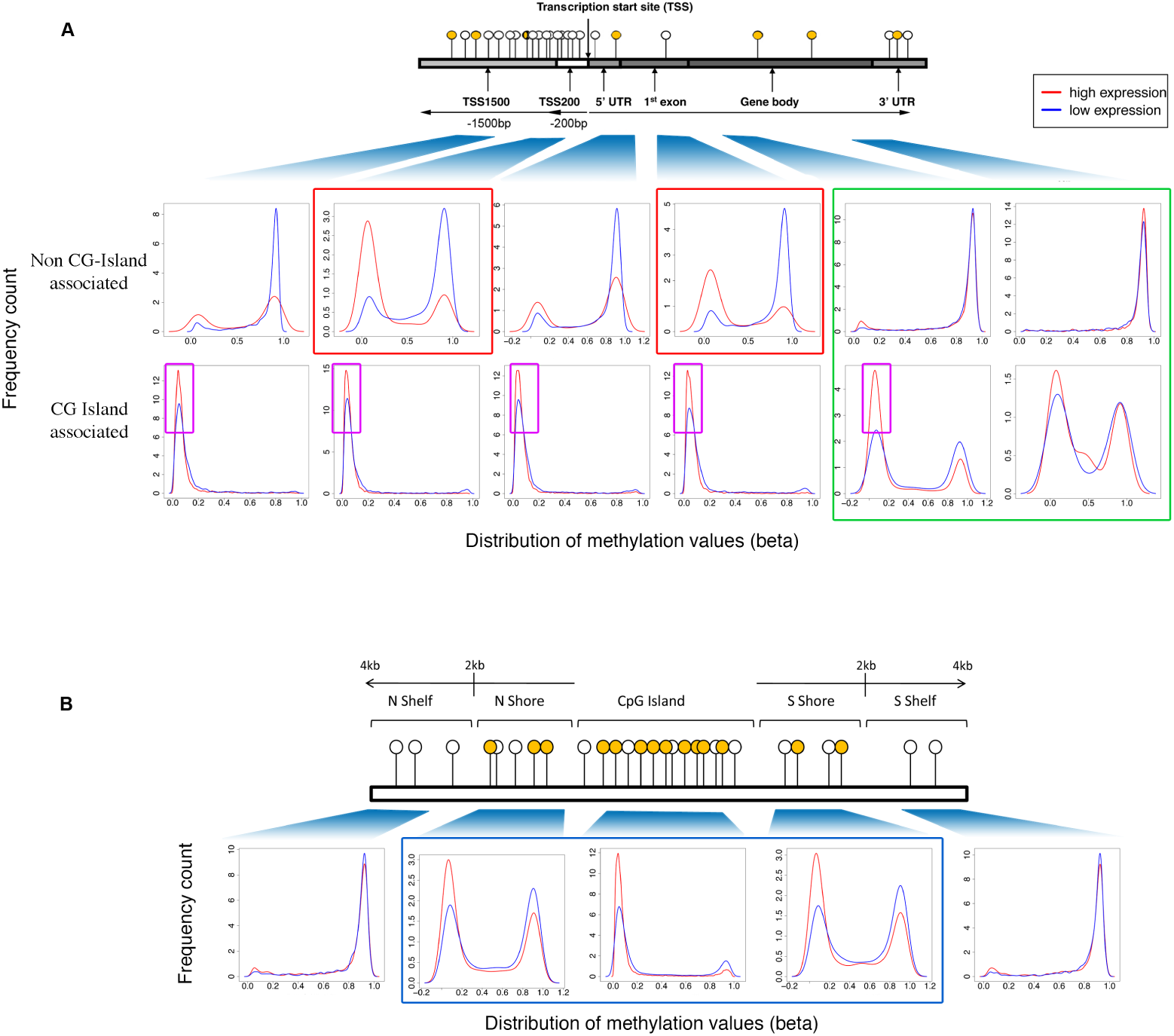
Relationship between DNA methylation and gene expression across regulatory features. (A) gene-associated genomic features (TSS1500, Tss200, 5’UTR, 1st exon, gene body, 3’UTR). Weighted density distributions of CpG methylation relative to gene expression status (high expression – red; low expression – blue) is shown stratified by the presence (A bottom panel) / absence (A top panel) of overlapping CpG island. Boxes in red indicate regulatory features exhibiting a classical gene silencing effect. Boxes in green show regulatory features that are generally highly methylated. Boxes in purple denote regions of hypomethylation (B) CpG island associated regions (CpG island, North and South Shore, North and South Shelf). Boxes in blue indicate regions where significant differences in the distributions were observed (P ¡ 0.05, Wilcoxon test).

Figure 2B shows the distribution of DNA methylation sites between high and low genes in CpG island associated regions. Formal testing revealed significant differences between of the distributions at islands and shores using the Wilcoxon test (P ¡ 0.05 for each region) between high versus low expressed genes. In agreement with several other previous findings,^8^ CpG islands were generally highly likely to be hypomethylated irrespective of gene expression status. In contrast, at gene-associated CpG island shores, DNA methylation and gene expression were generally anti-correlated, whilst CpG island shelves were consistently methylated for both high and low expressed genes.

We next examined whether any of the aforementioned patterns differ according to whether specific genes show evidence for dynamic regulation during development. We therefore performed a formal comparison of gene expression levels between birth and 12-month samples using a paired t-test and identified a total of 1, 362 genes (225 upregulated; 1137 downregulated) that change dynamically in CD4+ T-cells in early life (Adj. P. Value <0.05). HM450k probes associated with these genes were identified using the manufacturers annotations, and we compared their density distributions against the remaining stable genes (all genes minus 1137 dynamic genes) across several genomic regions. We found no general deviations in the overall relationship between methylation and gene expression between stable and dynamic genes. However We did observe a more pronounced shift in the methylation distribution between stable and dynamic genes in CpG islands and shores (figure 3B). We also noted a substantial difference in the level of methylation within the gene body in dynamic genes (figure 3C). In general terms these comparatively different distributions suggest that CpG islands, shores and the gene bodies exhibited substantial de-methylation during early development associated with changes in gene expression levels.

**Figure 3.**
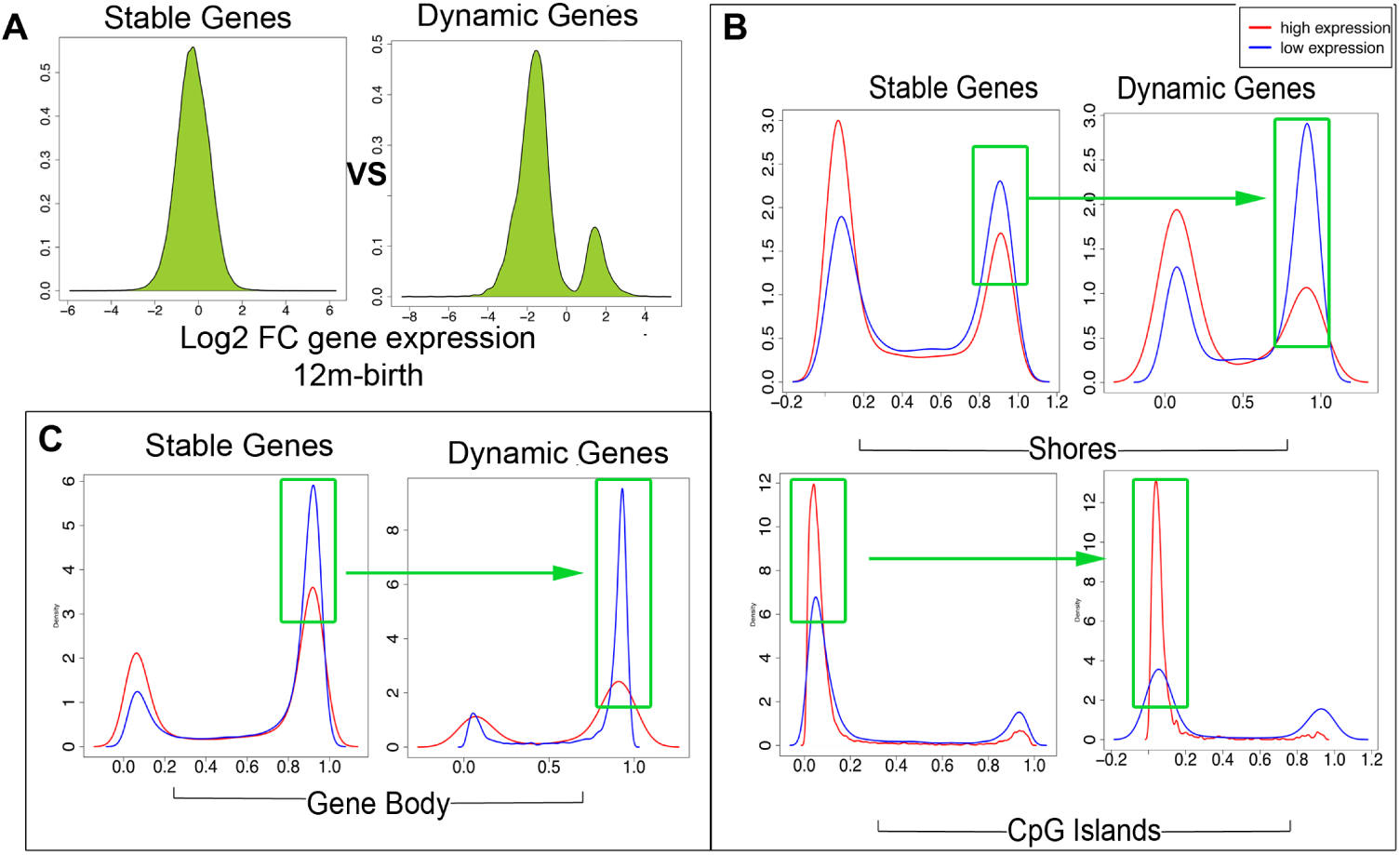
Dynamic versus stable gene expression is associated with variations in DNA methylation in CpG islands and shores and in the gene body. Dynamic genes were defined as those significantly (adj P less than 0.05, logFC 2) differentially expressed between 12-months and birth in the CD4+ T-cells. Stable genes were the remaining genes in the data set. The distribution of DNA methylation was plotted for high versus low expressed genes for stable genes, and compared with the distribution of DNA methylation between up and down regulated dynamic genes. Differences were noted in the CpG islands and shores (B) and in the gene body (C). Green boxes highlight differences in the area under the curve

## Discussion

One of the major challenges in Epigenome-Wide Association Studies (EWAS) is extrapolating likely functional consequences, in terms of likely effect of gene expression, of methylation changes associated with phenotypic/exposure groups of interest. In this study we have analysed the general relationship between gene expression and DNA methylation in CD4+ T-lymphocytes at two time-points using the widely utilized HM450K methylation platform in combination with array-based gene expression data. Although limited to around 2 percent of total CpG methylation sites in the genome, these cover more than 95 percent of all RefSeq genes and therefore allow general relationships with gene expression to be explored. It is important to note that we have not explored the role of enhancer elements and other distal regulatory DNA regions, widely recognized as playing a major role in gene regulation. Furthermore we have considered only CpG methylation, in only one cell type and thus our results should be interpreted within these limits. Nevertheless our we have identified some general associations of interest pertaining to the link between methylation and gene expression status using the HM450K platform. As anticipated we found that methylation in close proximity to the transcriptional start site (TSS200, 1st Exon) generally correlates strongly with variations in gene expression whilst distal regions further downstream of the TSS do not (gene body and 3’UTR). This likely reflects the differing roles of DNA methylation in these different contexts. It has been shown by others that DNA methylation at the TSS may block the initiation of transcription^3^,^4^ whilst methylation in the gene body may be more important for controlling gene splicing^9^ and control of developmental timing of expression,^10^ and our data supports this. It is possible that gene body methylation may be highly correlated with alternative transcript isoforms, and we were not able to address this in the present study. The CpG-island associated promoter regions were generally unmethylated for all genes as expected, and the level of de-methylation in these regions allows for fine-tuning of gene-expression levels. Such promoters are likely regulated by alternative epigenetic processes such as changes in the pattern of histone modification and polycomb repressive proteins.^11^ In contrast, substantial fluctuations in methylation levels in non-CpG island promoters were highly associated with variations in gene expression levels. This is consistent with the more recent findings regarding the role of methylation in non-CpG Island promoters^4^.^12^ The role of DNA methylation in the gene body and the 3’UTR is unclear, these regions showed little associations with gene expression levels and therefore expression status of underlying genes cannot generally be inferred from methylation data in isolation. Indeed methylation at the 3’UTR appears entirely independent from expression level. Interestingly we did observe a substantial gain in methylation in the gene bodies associated with the downregulation of gene expression in developmentally regulated genes. Others have previously shown that gene-body methylation varies with the rate of differentiation and cell division history of the tissue.^10^ Methylation in the gene-body may therefore be important in lineage commitment and control gene splicing and the utilization of alternative transcripts over this developmental period^9^.^13^ In the CpG island regions we found the island shores to be the most highly correlated with gene expression consistent with other reports^14^.^15^ Curiously the CpG island shelves were completely methylated for all genes tested using this platform, but the significance of this is unclear. In summary our findings confirm that DNA methylation is not always associated with gene silencing, highlighting the need to consider genomic context when interpreting epigenetic studies. For candidate prioritization for EWAS we recommend considering the proximity to the promoter and the presence of underlying CpG island as additional criteria, but reiterate the difficulty in inferring methylation status from cross sectional DNA methylation data in isolation.

## Methods

### Data sets

The data set analysed was published by us^7^ and available from the GEO repository (http://www.ncbi.nlm.nih.gov/geo/, GSE34639). Blood samples were collected from 48 individuals at birth and again at 12-months. Half the samples were either cultured in AIM-V media alone or with media containing anti-CD3 antibody and interleukin-2 for 24 hours; total CD4+ T-cells were then isolated by magnetic bead positive selection and DNA and total RNA were co-purified using the Qiagen Allprep kit (Qiagen). DNA from two individuals was pooled into a single sample at each time point, resulting in 12 conventional and 12 anti-CD3 birth samples, and 12 conventional and 12 anti-CD3 12 months samples. One microgram of genomic DNA samples were treated with sodium bisulphite (Human Genetic Signatures) according to manufacturers instruction and DNA methylation was measured on the Illumina Human Methylation 450k array (Illumina). Total RNA samples were also pooled from conventional and anti-CD3 cultures at each age. Gene expression was measured using Affymetrix Human Gene 1.0ST arrays. For the purposes of this analysis, we considered the conventional cells only as the anti-CD3 activation protocol did not induce a methylation response and has been published previously.^7^

### Data processing and analysis

All analysis was performed in R (www.cran.r-project.org) using analysis packages available through the bioconductor project (www.bioconductor.org). For methylation data, raw iDat files were imported into R using the Minfi package.^16^ Array preprocessing was performed using the Illumina method with normalization to controls. The preprocessed data underwent a within-array subset quantile normalization step^17^ to remove technical bias attributable to the different probe chemistries between Type1 and Type II probes. Any poor-performing probes with a detection P-value call ≥ 0.01 for 1 or samples were removed from the data set, as well as cross-hybridizing probes that provide unreliable data.^18^ Probes on the X and Y-chromosomes were removed prior to analysis to remove sex-specific effects. Beta values were derived from the processed data defined as the ratio of methylated probe intensity to the overall intensity given by *Beta* = *methylated/*(*methylated* + *umethylated*) * 100). Affymetrix .CEL files were imported into the Expression Console software (Affymetrix, Santa Clara) and preprocessed with PLIER (gcbc background correction, quantile normalization, iterPLIER summarization). Processed data were converted to the Log2 scale and variance stabilized by adding 16 to all data points. Affymetrix arrays were annotated using the hugene10sttranscriptcluster.db package to provide mappings between manufacturers probe IDs and ENTREZ gene IDs. Kernel density estimates were performed using the density function in the R base package with Gaussian smoothing kernel and default bandwidth settings, weighted by the number of observations in each class. To formally test differences in kernel density distributions we used the Wilcoxon signed rank test. To test for differential gene expression between birth and 12-months, an moderated t-test was performed using an empirical bayes procedure according to Smythe et al.^19^ Differential expression was defined as a greater than 2 fold change between birth and 12-months and false discovery rate adjusted P-value of less than 0.05.

### Region - gene associations

Illumina arrays were annotated using the IlluminaHumanMethylation450k.db package to provide mappings between manufacturers probe IDs and ENTREZ gene IDs. To categorize CpG sites into regulatory features (around the transcriptional start site) and CpG islands we used the annotations provided by the IlluminaHumanMethylation450k.db package. In cases where these regulatory features were not exclusive, i.e. a CpG may map to several features, in which case a count was given in each category.

## Author contributions statement

D.M performed the experiments and analyzed the results. R.S conceived the experiments. All authors reviewed the manuscript.

## Additional information

**Competing financial interests** The authors declare no competing financial interests.

The corresponding author is responsible for submitting a competing financial interests statement on behalf of all authors of the paper. This statement must be included in the submitted article file.

